# species2vec: A novel method for species representation

**DOI:** 10.1101/461996

**Authors:** Boyan Angelov

## Abstract

Word embeddings are omnipresent in Natural Language Processing (NLP) tasks. The same technology which defines words by their context can also define biological species. This study showcases this new method - species embedding (`species2vec`). By proximity sorting of 6761594 mammal observations from the whole world (2862 different species), we are able to create a training corpus for the skip-gram model. The resulting species embeddings are tested in an environmental classification task. The classifier performance confirms the utility of those embeddings in preserving the relationships between species, and also being representative of species consortia in an environment.

## Introduction

Species Distribution Models have historically been focused on the estimation of a potential ecological niche for a single species [5]. One of the largest drawbacks of this approach is that those models do not take into account any possible interactions between the organisms (which are, of course, commonplace in nature) [4]. Such interactions include mutualism, commensalism, predation, and symbiosis [7]. Thus there is a growing need for methods that include this valuable information. Such tools can improve our understanding of ecosystems and their communities and direct more efforts into conservation and assessment of climate change effects on the biosphere. Recent advances in other areas of machine learning, such as NLP open opportunities for cross-domain adoption. One technique that is very common in the industry is the generation of word embeddings [10]. The current study aims to implement this technique, prove its statistical and ecological significance, and provide the research community with a new concept for species and an associated tool to use. A similar project (but a different approach, focused on environmental data) was performed by Chen et. al [4], unfortunately, the code and model was not made publically available, so we could not compare the methods directly.

## Materials and Methods

The data processing pipeline is shown on Figure 1, where the steps are in sequential order.

**Figure 1.**
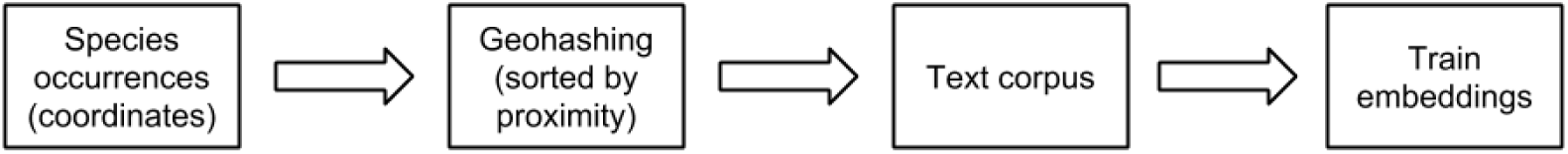
Data processing pipeline.

### Occurence data

Species occurence data was obtained from Global Biodiversity Information Facility (GBIF) [6]. The data for the species composition of different environments is obtained by a custom data download [9]. For initial version of `species2vec` the subject are mammal species (class Mammalia), because the ecological interpretation for testing is easier (there is more descriptive data for them available, as compared to other groups such as insects etc.). 6761594 records from 1760 published datasets were obtained. 2862 unique species were present in the dataset. In order to test that the generated `species2vec` embeddings can ecologically characterize an environment, they were used to substitude occurence counts data for several mammal species in three different environment types (based on a study by Michael et al. [9]).

### Geohashing

Geohashing is computed by using the algorithm developed by Niemyer [11]. This technique allows for the conversion of geographic coordinates (lattitude and longitude) into hash codes. Those are then sorted alphanumerically, ordering the coordinate points by proximity.

### Data processing and embeddings training

Word embedding is a method for generating a vector (”embedding”) for each word based on its context. There are several different approaches to generating those, most of them relying on shallow neural networks. For this study the python `fasttext` library from Facebook Research [2] was used to generate the embeddings. `fasttext` relies on the extension of the continious skip-gram model by using subword information. The resulting embeddings are the sums of the n-gram vectors. For accessing and storing the embeddings the `Gensim` library was used [12]. In order for complete species names to be accurately predicted, the empty spaces were replaced by underscores (i.e. Grammomys ibeanus → Grammomys_ibeanus). Those names were placed inside a large corpus, ordered by proximity, and finally fed to `fasttext`. The basic objective of the Skip-gram model is to maximize the average log probability shown in Equation 1, where *w*_1_, *w*_2_, *w*_3_…*w*_*T*_ are the training words and *c* is the training context.

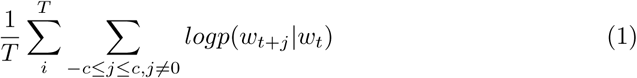

### Machine learning

Machine learning was performed by the `mlr` package [1]. The Random Forests algorithm was used because of its good out-of-the-box performance [3]. The dataset for model training was built by replacing the occurence counts for the several different mammal species by the generated embeddings. The vectors were weighted in order to account for the relative abundance (since it is not a presence-absence only dataset). A 300 × 5 matrix was created, split randomly into training and tests sets (50% each) and a model was trained. In order to visualise the embeddings, the t-Distributed Stochastic Neighbor Embedding (t-SNE) method was used [8].

## Results

### Species embeddings

After training the `species2vec` model based on the steps in Figure 1, *n* = 1987 unique species remain (69% of the original dataset). For the lost 31% not enough data points were available. The geohashing took ∼ 14 minutes on a 4 core 2.70GHz CPU 32GB RAM machine. The first step of investigating the embeddings performance is to use a function from the `Gensim` package which allows for the extraction of most similar species embeddings. The results are shown on Figure 2. The *Grammomys ibeanus* species was chosen for this, since it is one of the species in the environment dataset which is used to test the embeddings. All 10 most similar species are also rodents from central Africa.

**Figure 2.**
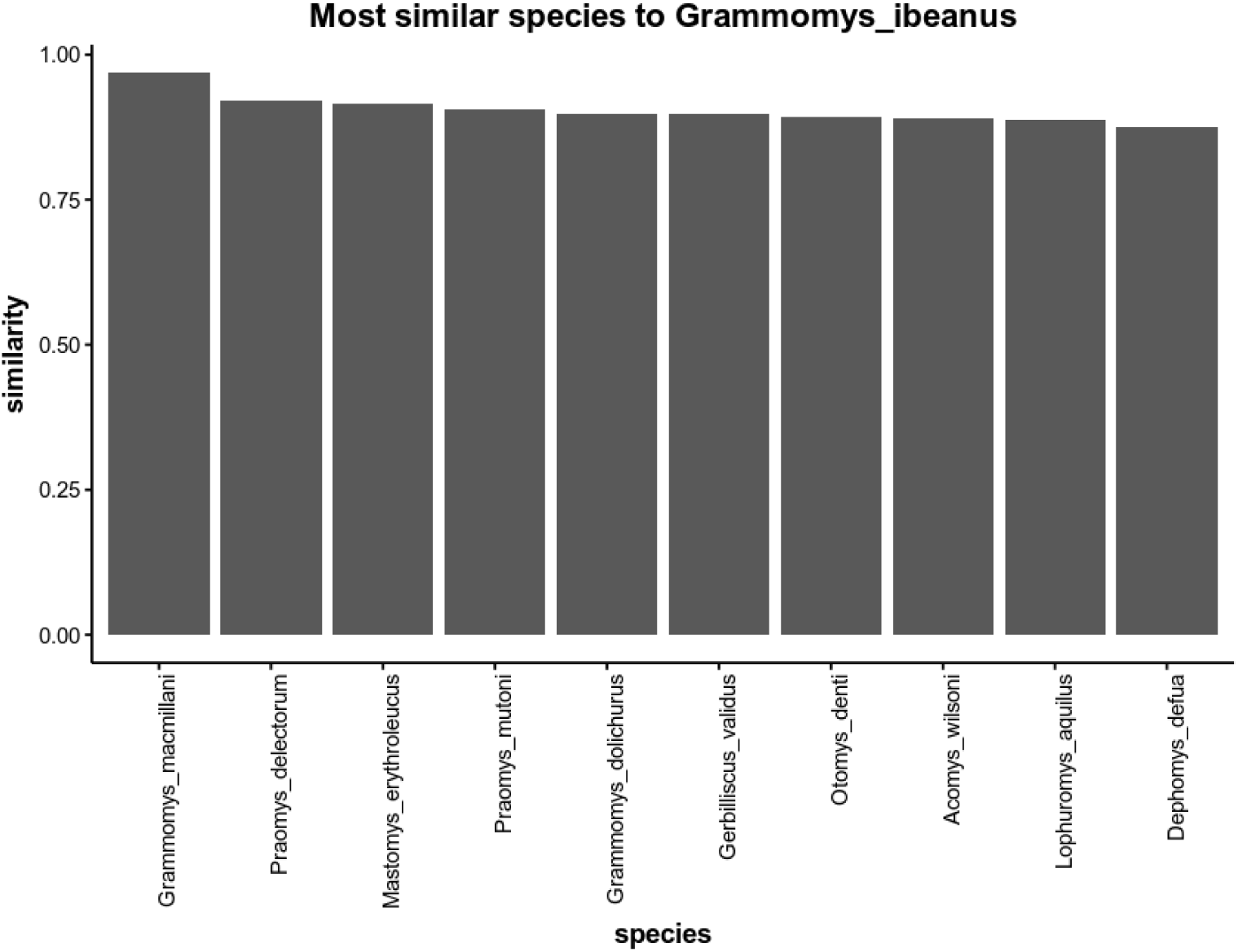
Most similar species. All shown species are central african rodents.

A second step in assessing the quality of the species embeddings is t-SNE visualisation. This result is shown on Figure 3. Samples of the data points are manually investigated to confirm that the clusters overlayed on top of the plot are correct. We can see that there is good separation between the continents, further confirming our hypothesis that the geohashing and subsequent embeddings have an ecological meaning. This large t-SNE plot has a lot of detail available for further interpretation. As an example we can zoom into one of the clusters to inspect the results deeper. This visualisation is shown on Figure 4. In the upper-left corner a mini cluster of the *Leontocebus genus* is present, showing that those several species of tamarins are closely related.

**Figure 3.**
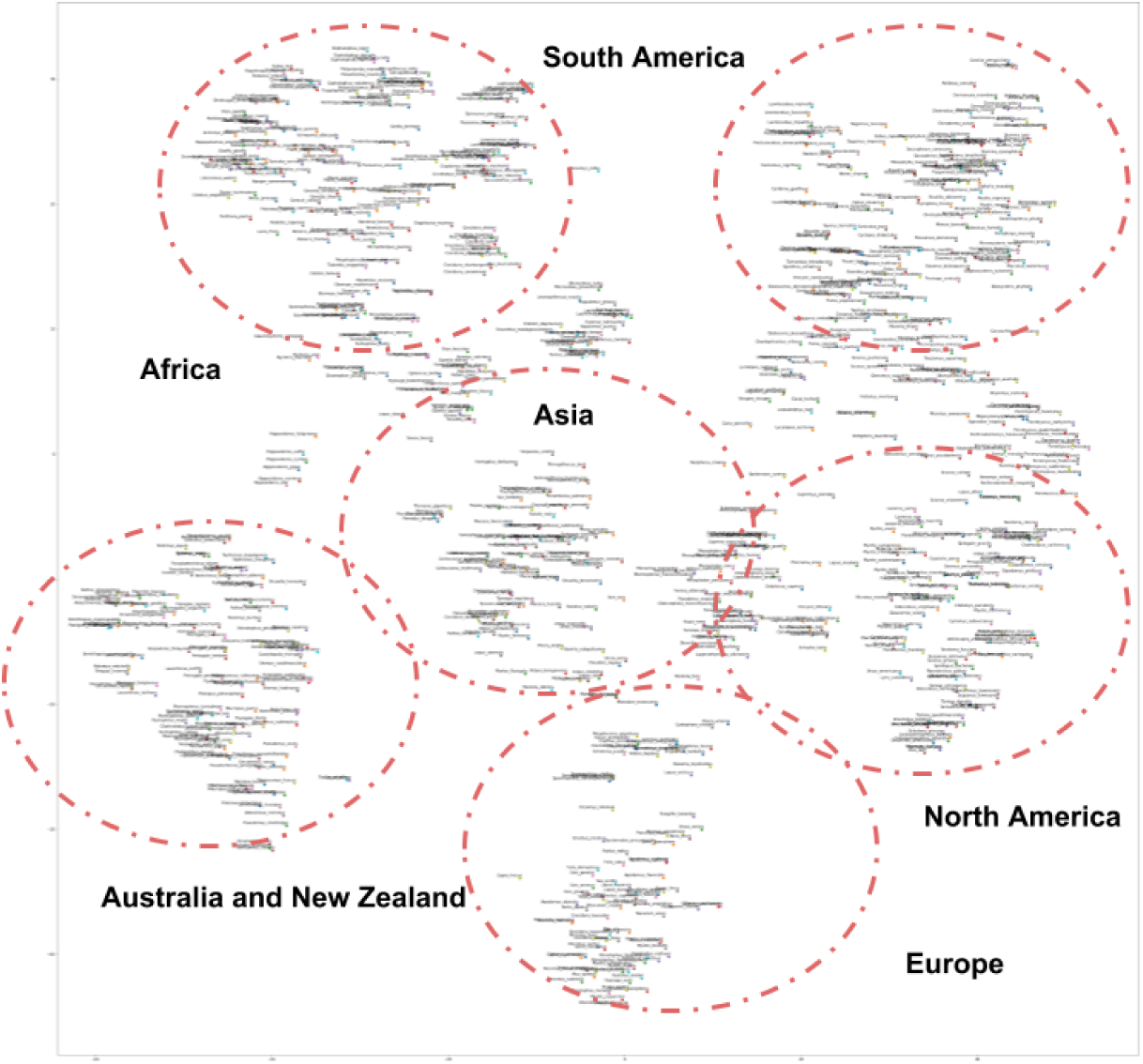
t-SNE plot of a sample of *n* = 1000 different embeddings. The red dotted lines denote the different clusters.

**Figure 4.**
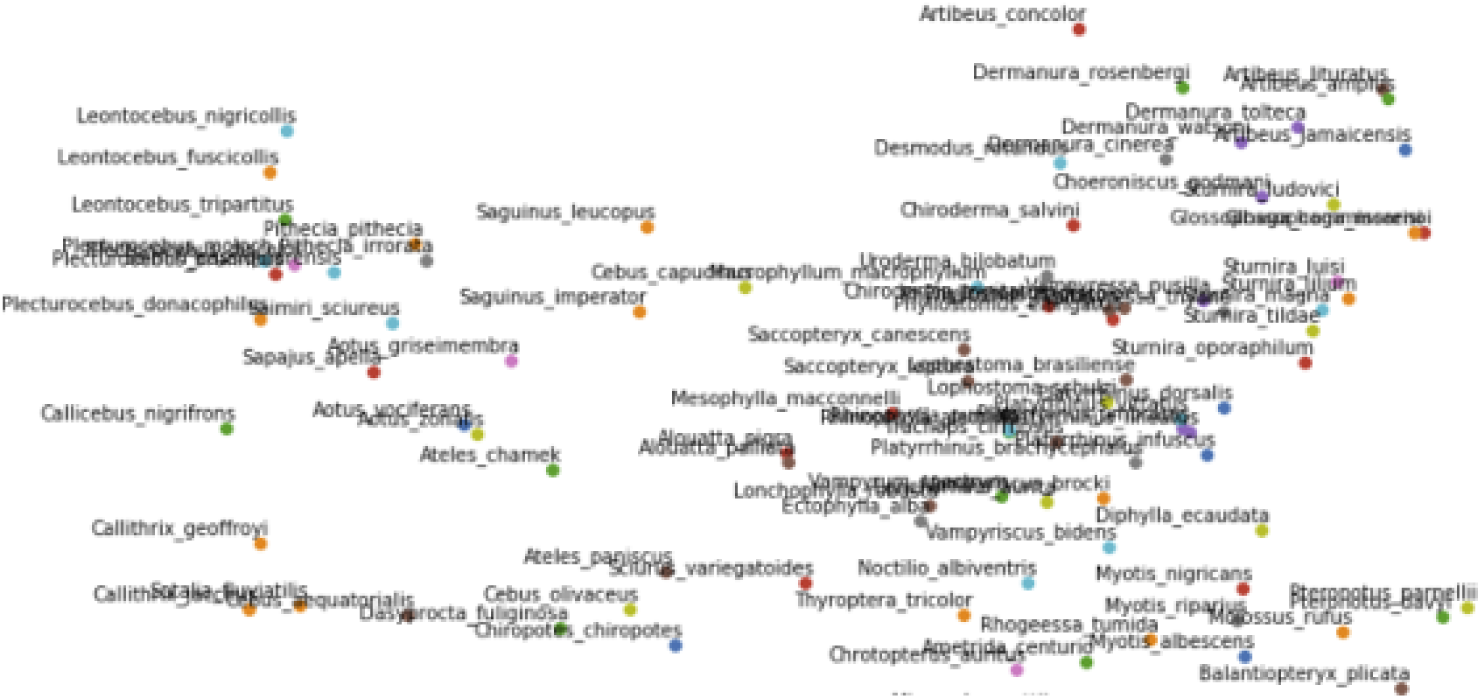
South American t-SNE cluster.

### Predicting environment types

The Random Forest classifier which was trained to predict the 3 different environments by using weighted `species2vec` embeddings achieved *accuracy* = 87% and *mmce* = 0.13^1^. The decision boundaries are plotted on Figure 5, showing a clear separation between the environment types^2^.

**Figure 5.**
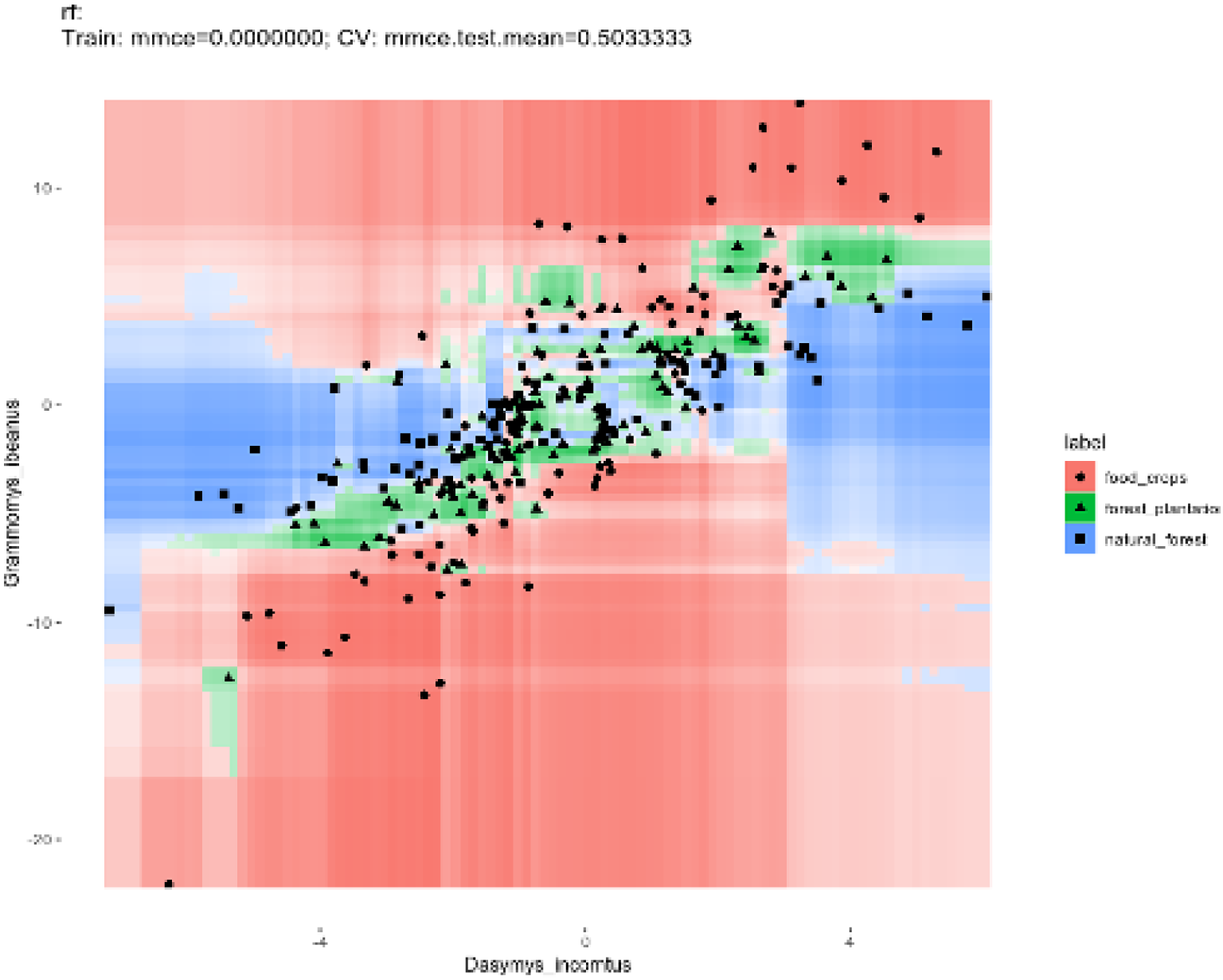
Classifier decision boundary.

## Discussion

The results obtained in this study prove that the concept of generating species embeddings is meaningful statistically and ecologically. The initial observation of the most similar species (Figure 2) confirms that the vector generated for *Grammomys ibeanus* indeed represents it as occupying the same niche as the other listed species (*Grammomys macmillani* ^3^, *Praomys delectorum, Mastomys erythroleucus* and others). The t-SNE visualisation (Figure 3) provides further evidence for this. Some information is lost by reducing the dimensions, but still the separation between different biotopes is clear. A potential future improvement of this plot is to generate an interactive 3D visualisation that would allow for even deeper exploration, since even more deeper patterns in local communities can be discovered (i.e. can we see a separation between sub-saharan african species and those to the north?). The high classification accuracy for the environment type classification dataset proves that a vector representation of a species can also represent its environment successfully. Environments represented by the weighted associated species vectors could be distunguished strongly, even with a low number of features (≤ 6) and having data from relatively similar environments.

`species2vec` as a method can be used in solving a variety of ecological problems and challenges. Domain experts in the field can benefit greatly from having a tool which can recommend which species are most likely to be encountered next in the environment. Related to this, researchers can assess and classify differences in habitats in a more accurate way by using the weighted vectorized species representations instead of simple raw counts. Those differences in habitats can have large ecological significance if they are caused by direct habitat descruction or climate change. Invasive species can also be classified by using their embeddings.

### Future work

The embeddings are available in a GitHub repository (https://github.com/boyanangelov/species2vec/) and also archived on Zenodo (https://zenodo.org/record/1477661). The current version of `species2vec` (v1.0) provides embeddings for mammals only. It can be interesting and useful to generate embeddings of other separate groups of organisms (insects, birds) or to include all taxonomic groups available in GBIF (also flora). The aforementioned interactive exploration tool can also prove to be useful to domain experts.

## Acknowledgments

The author wants to ackwowledge GBIF for providing the dataset used for the study.

## Footnotes

1 Mean missclassification error.

2 Here two species (features) plotted against each other.

3 Note that there are two species from the same genus in the list.

